# Potential T-cell and B-cell Epitopes of 2019-nCoV

**DOI:** 10.1101/2020.02.19.955484

**Authors:** Ethan Fast, Russ B. Altman, Binbin Chen

**Affiliations:** Nash, Menlo Park, CA, USA; Department of Bioengineering, Stanford University, Stanford, CA, USA; Department of Genetics, Stanford University, Stanford, CA, USA

**Keywords:** 2019-nCoV, COVID-19, Vaccine, Epitope, Antibody

## Abstract

As of early March, 2019-nCoV has infected more than one hundred thousand people and claimed thousands of lives. 2019-nCoV is a novel form of coronavirus that causes COVID-19 and has high similarity with SARS-CoV. No approved vaccine yet exists for any form of coronavirus. Here we use computational tools from structural biology and machine learning to identify 2019-nCoV T-cell and B-cell epitopes based on viral protein antigen presentation and antibody binding properties. These epitopes can be used to develop more effective vaccines and identify neutralizing antibodies. We identified 405 viral peptides with good antigen presentation scores for both human MHC-I and MHC-II alleles, and two potential neutralizing B-cell epitopes near the 2019-nCoV spike protein receptor binding domain (440-460 and 494-506). Analyzing mutation profiles of 68 viral genomes from four continents, we identified 96 coding-change mutations. These mutations are more likely to occur in regions with good MHC-I presentation scores (p=0.02). No mutations are present near the spike protein receptor binding domain. Based on these findings, the spike protein is likely immunogenic and a potential vaccine candidate. We validated our computational pipeline with SARS-CoV experimental data.

**Significance Statement:** The novel coronavirus 2019-nCoV has affected more than 100 countries and continues to spread. There is an immediate need for effective vaccines that contain antigens which trigger responses from human T-cells and B-cells (known as epitopes). Here we identify potential T-cell epitopes through an analysis of human antigen presentation, as well as B-cell epitopes through an analysis of protein structure. We identify a list of top candidates, including an epitope located on 2019-nCoV spike protein that potentially triggers both T-cell and B-cell responses. Analyzing 68 samples, we observe that viral mutations are more likely to happen in regions with strong antigen presentation, a potential form of immune evasion. Our computational pipeline is validated with experimental data from SARS-CoV.

---

**C**oronaviruses (CoV) first became widely known after the emergence of severe acute respiratory syndrome (SARS) in 2002. They are are positive, single-strand RNA viruses that often infect birds and a variety of mammals and sometimes migrate to humans (1). While coronaviruses that infect humans usually present mild symptoms (e.g. 15% of cases involving the common cold are caused by a coronavirus), several previous outbreaks, such as SARS and Middle East respiratory syndrome (MERS) (2), have caused significant fatalities in infected populations (3).

In December 2019, several patients with a connection in Wuhan, China developed symptoms of viral pneumonia. Sequencing of viral RNA determined that these cases were caused by a novel coronavirus named 2019-nCoV or SARS-CoV-2 (4, 5). The virus has infected more than 100,000 people across 100 countries as of March 7th 2020 according to a World Health Organization report (6). COVID-19, the disease caused by 2019-nCoV, has led to the deaths of thousands of patients and is clearly transmissible between humans (3).

The deployment of a vaccine against 2019-nCoV would arrest its infection rate and help protect vulnerable populations. However, no vaccine has yet passed beyond clinical trails for any coronavirus, whether in humans or other animals (7, 8). In animal models, inactivated whole virus vaccines without focused immune epitopes offer incomplete protection (8). Vaccines for SARS-CoV and MERS-CoV have also resulted in hypersensitivity and immunopathologic responses in mice (7, 9). The diversity among coronavirus strains presents another challenge (1, 9), as there are at least 6 distinct sub-groups in the coronavirus genus. And while SARS-CoV and 2019-nCoV share the same subgroup (2b), their genomes are only 77% identical (4). Understanding the shared and unique epitopes of 2019-nCoV will help the field design better vaccines or diagnostic tests for patients.

Both humoral immunity provided by B-cell antibodies and cellular immunity provided by T-cells are essential for effective vaccines (10, 11). Though humans can usually mount an antibody response against viruses, only neutralizing antibodies can fully block the entry of viruses into human cells (12). The location of antibody binding sites on a viral protein strongly affects the body’s ability to produce neutralizing antibodies; for example, HIV has buried surface proteins that protect themselves from human antibodies (13). For this reason, it would be valuable to understand whether 2019-nCoV has potential antibody binding sites (B-cell epitopes) near their interacting surface with its known human entry receptor Angiotensin-Converting Enzyme 2 (ACE2).

Beyond neutralizing antibodies, human bodies also rely upon cytotoxic CD8 T-cells and helper CD4 T-cells to fully clear viruses. The presentation of viral peptides by human Major Histocompatibility Complex (MHC or HLA) class I and class II is essential for anti-viral T-cell responses (14). In contrast to B-cell epitopes, T-cell epitopes can be located anywhere in a viral protein since human cells can process and present both intracellular and extracellular viral peptides (15).

These immunology principles guide the design of our current study (Fig. 1). First we focus on T-cell epitopes. 2019-nCoV carries 4 major structural proteins—spike (S), membrane (M), envelope (E) and nucleocapsid (N)—and at least 6 other open reading frames (ORFs) (1, 16). All protein fragments have the potential to be presented by MHC-I or MHC-II and recognized by T-cells. We apply NetMHCpan4 (17) and MARIA (15), two artificial neural network algorithms, to predict antigen presentation and identify potential T-cell epitopes.

**Fig. 1.**
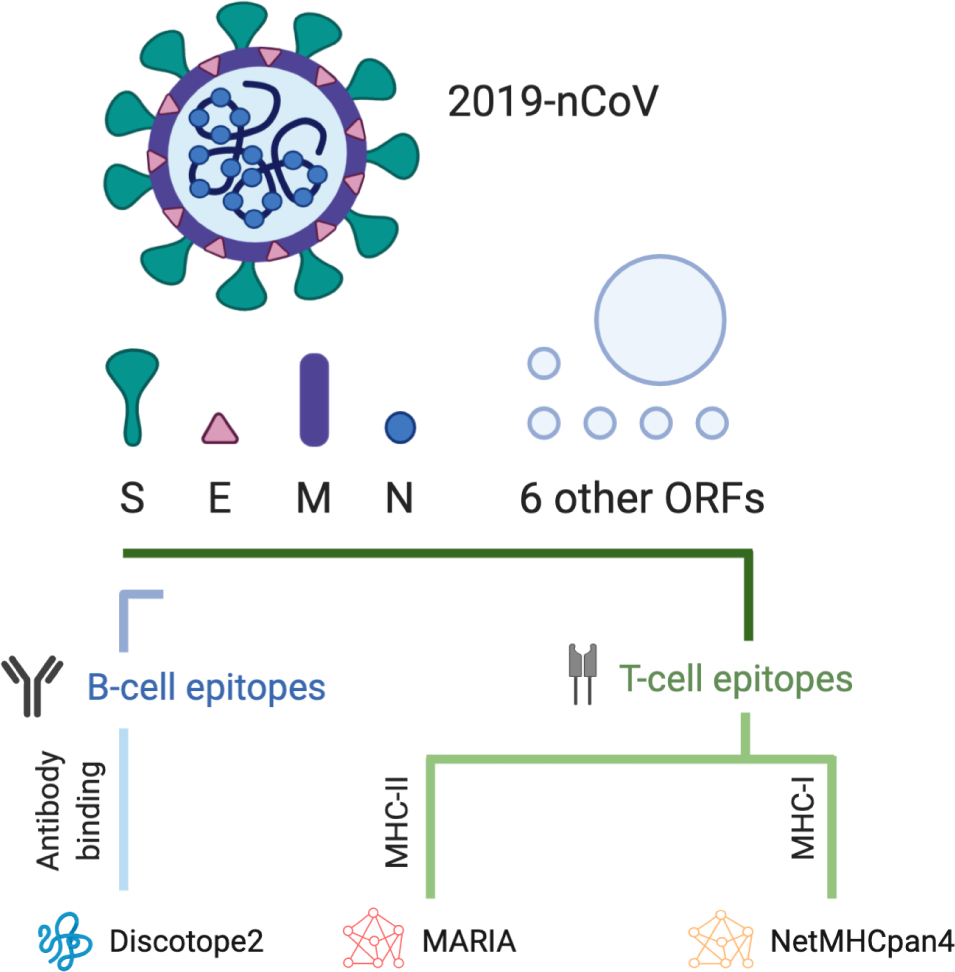
Computational pipeline to identify epitopes in 2019-nCoV. The 2019-nCoV genome codes for Spike (S), Membrane (M), Envelope (E), Nucleocapsid (N) and at least 6 other open reading frames (ORFs). The S protein is a target for antibodies and we model its 3D structure with Discotope2 to identify likely binding sites (B-cell epitopes). We scan peptide sequences from individual viral proteins with NetMHCPan4 and MARIA for MHC-I and MHC-II presentation to identify potential T-cell epitopes. Our MHC-I analysis include common alleles for HLA-A, HLA-B, and HLA-C, and our MHC-II analysis include common alleles of HLA-DR.

Next we focus on B-cell epitopes. The spike (S) protein is the main trans-membrane glycoprotein expressed on the surface of 2019-nCoV and is responsible for receptor binding and virion entry to cells (1, 18). We narrow our B-cell epitope search to S protein since it is the most likely target of human neutralizing antibodies. We first perform homology modeling to estimate the full 2019-nCoV spike protein structure and compare our results to partially solved spike protein structures (18–20). Discopte2 uses a combination of amino acid sequences and protein surface properties to predict antibody binding sites given a protein structure (21). We use this tool to identify potential 2019-nCoV B-cell epitopes as the co-crystal structure of 2019-nCoV’s spike protein and antibody is not yet available.

Finally, we examine the viral mutation pattern. Global sequencing efforts (22) have enabled us to obtain a cohort of 68 2019-nCoV genomes from four continents. Based on the resulting viral mutation profile, we examine whether mutations are driven by the immune selection pressure. Highly mutated regions might be excluded from the vaccine candidate pool.

## Results

### T-cell epitopes of 2019-nCoV based on MHC presentation

We scanned 11 2019-nCoV genes with NetMHCpan4 and MARIA to identify regions highly presentable by HLA complexes (human MHC). A epitope is labeled positive if presentable by more than one third of common alleles among the Chinese population. We display a summary of our T-cell epitope findings in Table 1. In Fig. 2 we show regions of strong MHC presentation across HLA-A, HLA-B, HLA-C, and HLA-DR alleles, both at a medium threshold (Fig. 2A), and at a high threshold (Fig. 2B). Consistent with the similar finding from SARS-CoV, proteins ORF1AB, S, and E contain high numbers of presentable antigen sequences across both MHC-I and MHC-II (8). These highly presentable peptides on functional viral proteins provide a candidate pool for T-cell epitope identifications and epitope-focused vaccine design. The presentation scores of individual antigen peptides can be found in SI Appendix, Supplementary Tables 5 and 6.

**Table 1.** Summary of potential 2019-nCoV T-cell epitope candidates based on HLA antigen presentation scores. The HLA-A, HLA-B, HLA-C, and HLA-DR columns refer to the number of peptide sequences that were predicted to present across more than one third of common alleles among the Chinese population. Length refers to the number of amino acids in a given gene. HLA-C are more likely to present 2019-nCoV antigens across one third of alleles compared to HLA-A or B. Fisher’s exact test P-value levels are indicated with *** (< 0.001), ** (< 0.01), and * (< 0.05).

| Gene | Length | HLA-A |  | HLA-B |  | HLA-C |  | HLA-DR |  |
| --- | --- | --- | --- | --- | --- | --- | --- | --- | --- |
| ORF1ab | 7096 | 370 | 5.21% | 516 | 7.27% | 775*** | 10.9% | 918 | 12.9% |
| S | 1273 | 59 | 4.63% | 91 | 7.15% | 132*** | 10.4% | 95 | 7.46% |
| ORF3a | 275 | 23 | 8.36% | 16 | 5.82% | 34** | 12.4% | 11 | 4.00% |
| E | 75 | 11 | 14.7% | 8 | 10.7% | 12 | 16.0% | 3 | 4.00% |
| M | 222 | 12 | 5.41% | 18 | 8.11% | 27* | 12.2% | 14 | 6.31% |
| ORF6 | 61 | 6 | 9.84% | 5 | 8.20% | 8 | 13.1% | 1 | 1.64% |
| ORF7a | 121 | 7 | 5.79% | 8 | 6.61% | 11 | 9.09% | 3 | 2.48% |
| ORF7b | 43 | 4 | 9.30% | 2 | 4.65% | 3 | 6.98% | 0 | 0.00% |
| ORF8 | 121 | 6 | 4.96% | 6 | 4.96% | 9 | 7.44% | 2 | 1.65% |
| N | 419 | 9 | 2.15% | 19 | 4.53% | 24 | 5.73% | 24 | 5.73% |
| ORF10 | 38 | 2 | 5.26% | 3 | 7.89% | 8 | 21.1% | 1 | 2.63% |

**Fig. 2.**
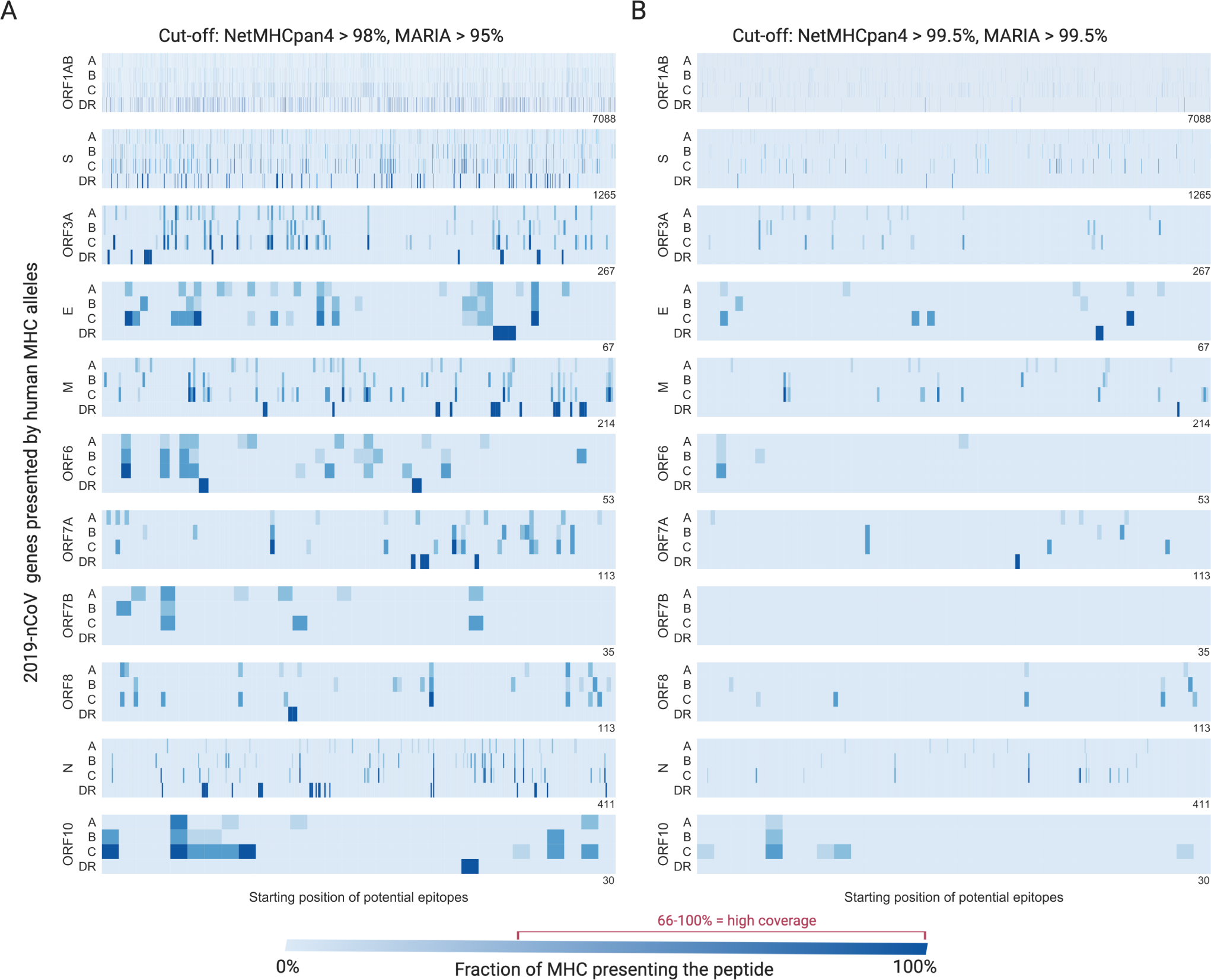
MHC presentation regions for 11 2019-nCoV genes across HLA-A, HLA-B, HLA-C, and HLA-DR. We plot results at both medium (**A**) and high (**B**) cut-offs for presentation. The X-axis indicates the position of each viral protein, and the y-axis indicates each human HLA gene family. Each blue stripe indicates the fraction of HLA alleles that can present a 9mer or 15mer viral peptide starting from a given position. A peptide being presented by more than one third of common alleles (red) is considered to be high coverage.

NetMHCpan4 predicts that a larger number of 2019-nCoV protein sequences can be presented by HLA-C alleles across all genes (Fisher’s exact test, *p* = 4.2*e*^−59^) compared to HLA-A or HLA-B (Table 1). Specifically, ORF1ab (*p* = 6.7*e*^−54^), S (*p* = 1.4*e*^−9^), ORF3a (*p* = 0.01), and M (*p* = 0.025) can be better presented by HLA-C alleles after Bonferroni correction.

### T-cell epitope validation

To estimate the ability of antigen presentation scores to identify 2019-nCoV T-cell epitopes, we performed a validation study with known SARS-CoV CD8 and CD4 T-cell epitopes (Fig. 3). From 4 independent experimental studies (23–26), we identified known CD8 T-cell epitopes (MHC-I, n=17) and known CD4 T-cell epitopes (MHC-II, n=3). We also obtained known non-epitopes (n=1236 and 246) for CD4, while for CD8 we include all 9mer sliding windows of S protein without a known epitope as non-epitopes. MHC-I presentation (with recommended cut-off 98% (17)) has a sensitivity of 82.3%, specificity of 97.0% and an AUC of 0.98 (Fig. 3A). MHC-II presentation (recommended cut-off 95% (15)) has a sensitivity of 66.6%, specificity of 91.1% and an AUC of 0.83 (Fig. 3B).

**Fig. 3.**
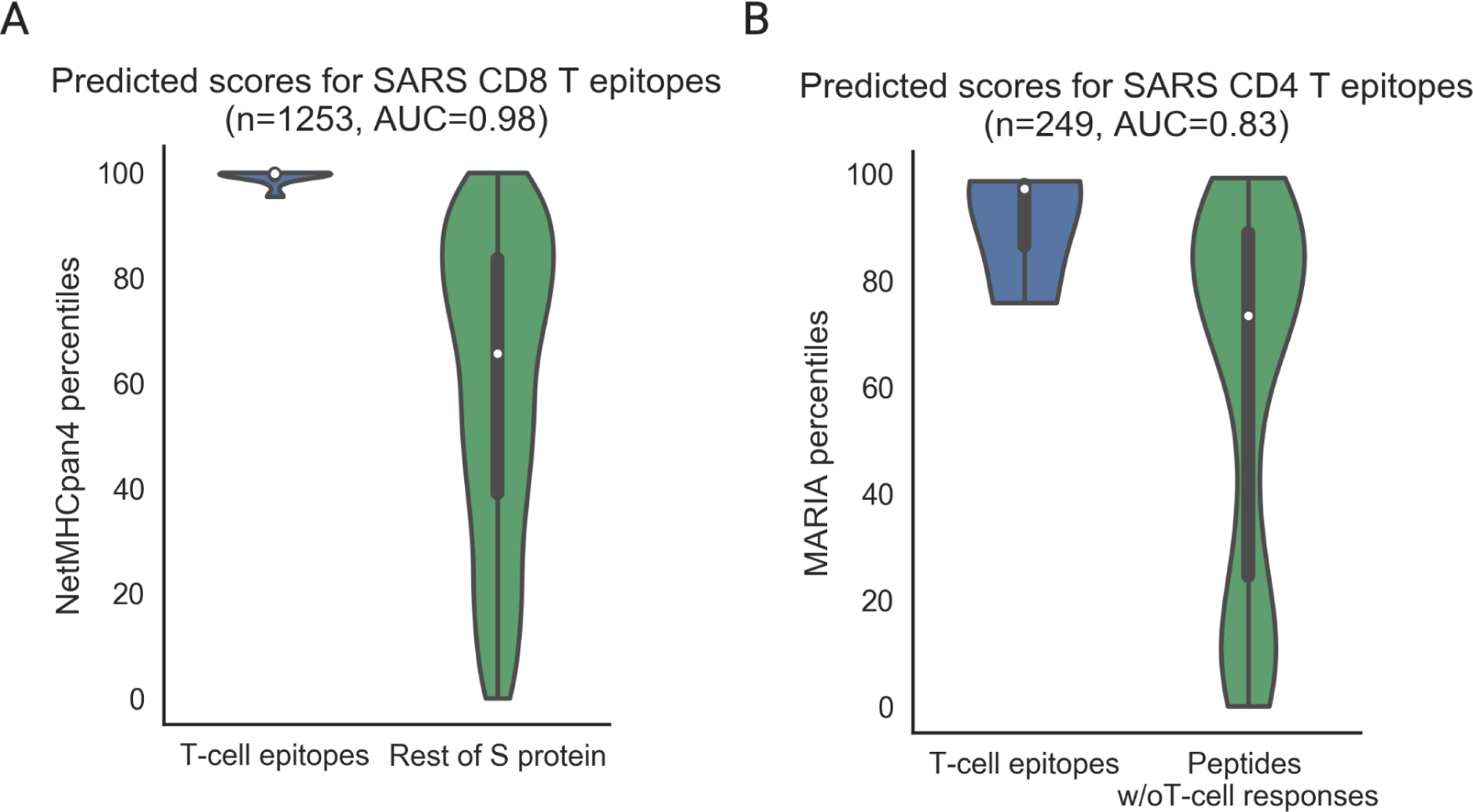
Validation of NetMHCpan and MARIA for T-cell epitope identification with SARS-CoV S protein epitopes. NetMHCpan4 and MARIA presentation percentiles were used to differentiate experimentally identified CD8 T-cell epitopes (MHC-I, n=17)(23–25) and CD4 T-cell epitopes (MHC-II, n=3)(26) from negative peptides (n=1236 and 246). Negative peptides for CD8 T-cell epitopes are all 9mer sliding windows of S protein without a reported epitope. Negative peptides for CD4 T-cell epitopes are experimentally tested. (**A)** NetMHCpan4 has a sensitivity of 82.3% and specificity of 97.0% with a 98th presentation percentile cut-off, and an overall AUC of 0.98. (**B)** MARIA has a sensitivity of 66.6% and specificity of 91.1% with a 95th presentation percentile cut-off, and an overall AUC of 0.83.

Given the high similarity between SARS-CoV and 2019-nCoV proteins, we expect our analysis on 2019-nCoV to perform similarly against future experimentally validated T-cell epitopes. The detailed scores and sequences of this analysis can be found in SI Appendix, Supplementary Tables 7 and 8.

### Viral mutation and antigen presentation

Human immune selection pressure has been shown to drive viral mutations which evade immune surveillance (e.g. low MHC presentation). We hypothesize that a similar phenomenon can occur in 2019-nCoV. We curated a cohort of 68 viral genomes across four continents and identified 93 point mutations, 2 nonsense mutation and 1 deletion mutation compared to the published reference genome (5). We plot point mutations against regions of MHC-I or MHC-II presentation in Fig. 4. The full protein sequence and mutation information can be found in SI Appendix, Dataset S2 and S3.

**Fig. 4.**
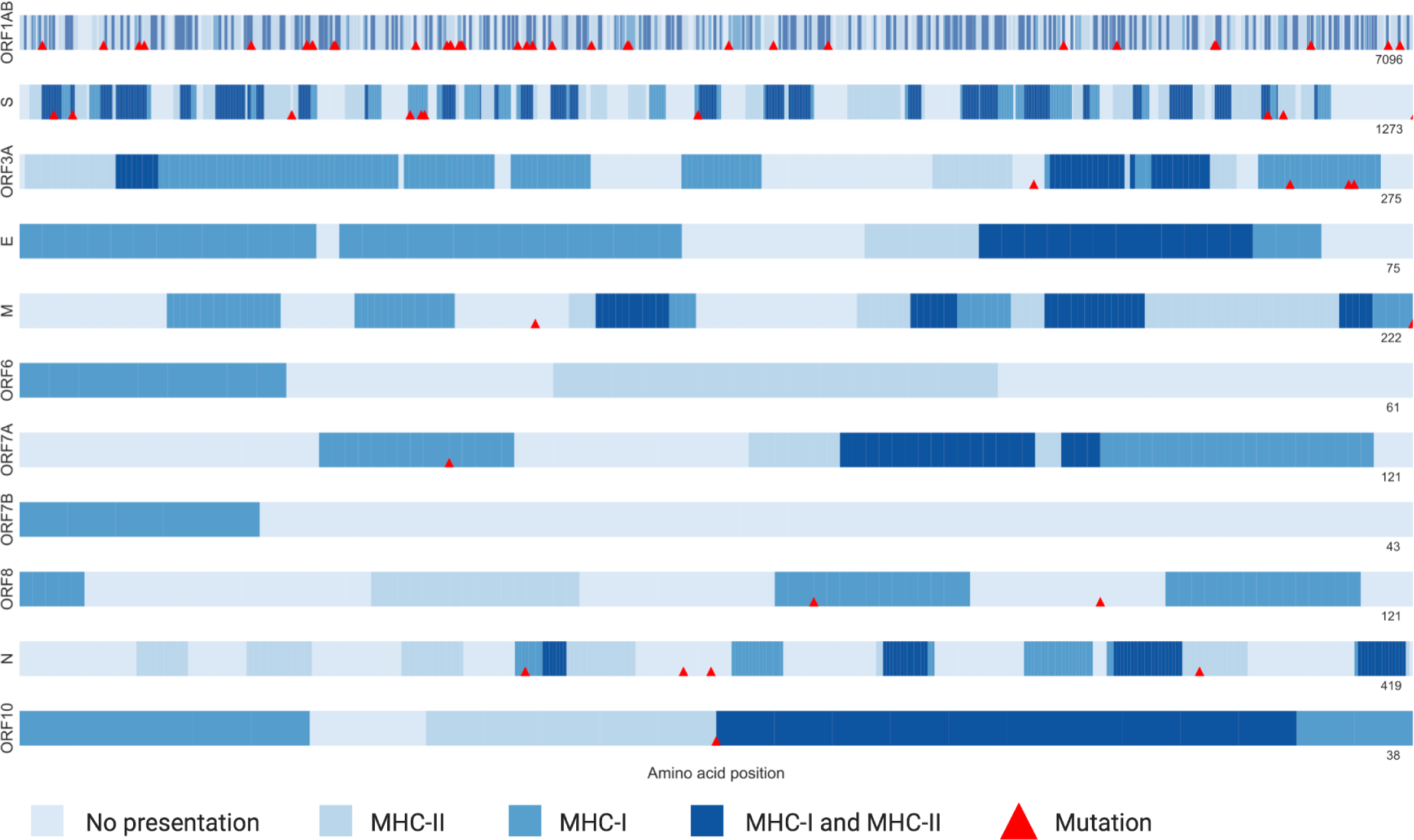
2019-nCoV point mutations correlates with MHC-I presentation regions. 93 point mutations at 59 unique positions (red triangles) are present in a cohort of 68 2019-nCoV cohort. Each row represents a viral protein and the shades of blue indicate whether MHC-I or MHC-II or both can present an antigen from this region. A region is labeled as presented by MHC-I or MHC-II if more than one third of the corresponding alleles can present the peptide sequences. Mutations are more likely be located in regions with MHC-I presentation (*p* = 0.02). No relationship was observed with MHC-II (*p* = 0.58) or MHC-I and MHC-II double positive region (dark blue, *p* = 0.14).

Mutations occur in most genes with the exception of E, ORF6, ORF7b, and N. We identify a recurrent mutation L84S in ORF8 in 20 samples. Mutations are more likely to occur in regions with MHC-I presentation (Fischer’s exact test, *p* = 0.02). No relationship was observed with MHC-II (*p* = 0.58) or an aggregate of MHC-I and MHC-II (*p* = 0.14). This is consistent with the role of CD8 T-cell in clearing infected cells and mutations supporting the evasion of immune surveillance (27).

We identified two nonsense mutations (ORF1AB T15372A and ORF7A C184T) in two viral samples from Shenzhen, China (EPI_ISL_406592 and EPI_ISL_406594). ORF7A C184U causes an immediate stop gain and a truncation of the last 60 amino acids from the ORF7A gene, which can be presented by multiple MHC alleles as indicated in our heatmap (Fig. 4). In a sample from Japan (EPI_ISL_407084), the viral gene ORF1AB loses nucleotides 94-118, which code for GDSVEEVL. An associated antigen FGDSVEEVL is predicted to be good presented by multiple MHC-I alleles (e.g. 99.9% for HLA-C*08:01, 99.2% for HLA-C*01:02, and 99.0% for HLA-B*39:01).

In summary, we observed that 2019-nCoV mutations often occur among or delete presentable peptides, a potential form of immune evasion. Patient MHC allele information and T-cell profiling are needed to better assess this phenomenon. However, patients typically carry fewer than 3 mutations, so a vaccine against multiple epitopes will not be nullified by such a small number of mutations.

### Potential B-cell epitopes of 2019-nCoV

We obtained the likely 3D structure of the full 2019-nCoV’s spike protein (SI Appendix, Dataset S4 and S5) by homology modeling with SARS-CoV’s spike protein (31). Given our understanding of SARS (32), the spike protein has two major conformations: up and down. The down position is more stable but less accessible to human entry receptors (e.g. ACE2). The receptor-binding domain (RBD) of the protein undergoes hinge-like conformational change to enter the up conformation, which is less stable but more accessible to human receptors (31). We compared our homology modeled structures to a later solved Cryo-EM structure (18) and found high similarity (Appendix SI, Fig. S1). Notably, the Cryo-EM solved structure has missing residues in the RBD due to limited electron density. Using modeled structures overcomes this issue.

We scanned spike protein structures with Discotope2 to identify potential antibody binding sites on the protein surface (Figs. 5 and 6). For the SARS-CoV S protein down conformation, we identified a strong cluster of antibody sites (>30 residues, pink or red) on the receptor binding domain (RBD, the ACE2 interacting surface). Three independent studies (28–30) discovered B-cell epitopes in this region (blue) with a combination of in vitro and ex vivo approaches. Additionally, Discotope2 identifies residue 541-555 as another binding site, which was supported by two independent studies (Fig. 5A, blue) (29, 30). This gives us some confidence in the ability of Discotope2 to predict B-cell epitopes if given a unknown protein structure.

**Fig. 5.**
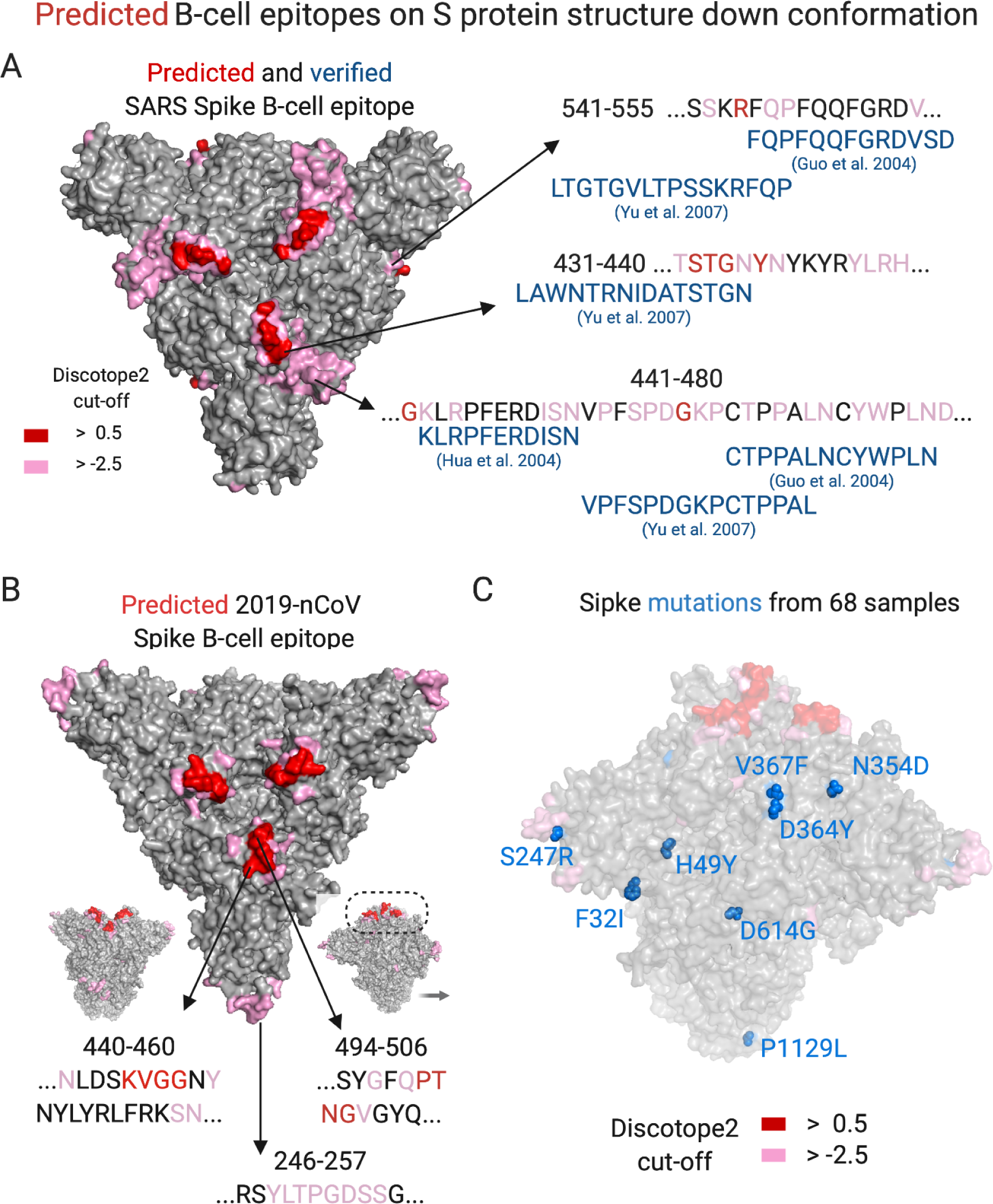
Predicted B-cell epitopes on SARS-CoV and 2019-nCoV spike (S) protein “down” position. 3D structures of both spike proteins in “down” position were scanned with Discotope2 to assess potential antibody binding sites (B-cell epitopes). Red indicates residues with score > 0.5 (cut-off for specificity of 90%), and pink indicates residues with score > −2.5 (cut-off for specificity of 80%). (**A**) The receptor binding domain (RBD) of SARS spike protein has concentrated high score residues and is comprised of two linear epitopes. One additional binding site is predicted to be around residue 541-555. Three independent experimental studies (28–30) identify patient antibodies recognize these three linear epitopes (blue). (**B**) Modeled 2019-nCoV spike protein is predicted to have a similar strong antibody binding site near RBD and an additional site around residue 246-257. (**C**) The predicted antibody binding site on 2019-nCov RBD potentially overlaps with the interacting surface for the known human entry receptor ACE2 (yellow). (**D**) Eight 2019-nCoV spike protein point mutations are present in a cohort of 68 viral samples. No mutations are near the ACE2 interacting surface. One mutation (S247R) occurs near one of the predicted antibody binding sites (residue 246-257).

**Fig. 6.**
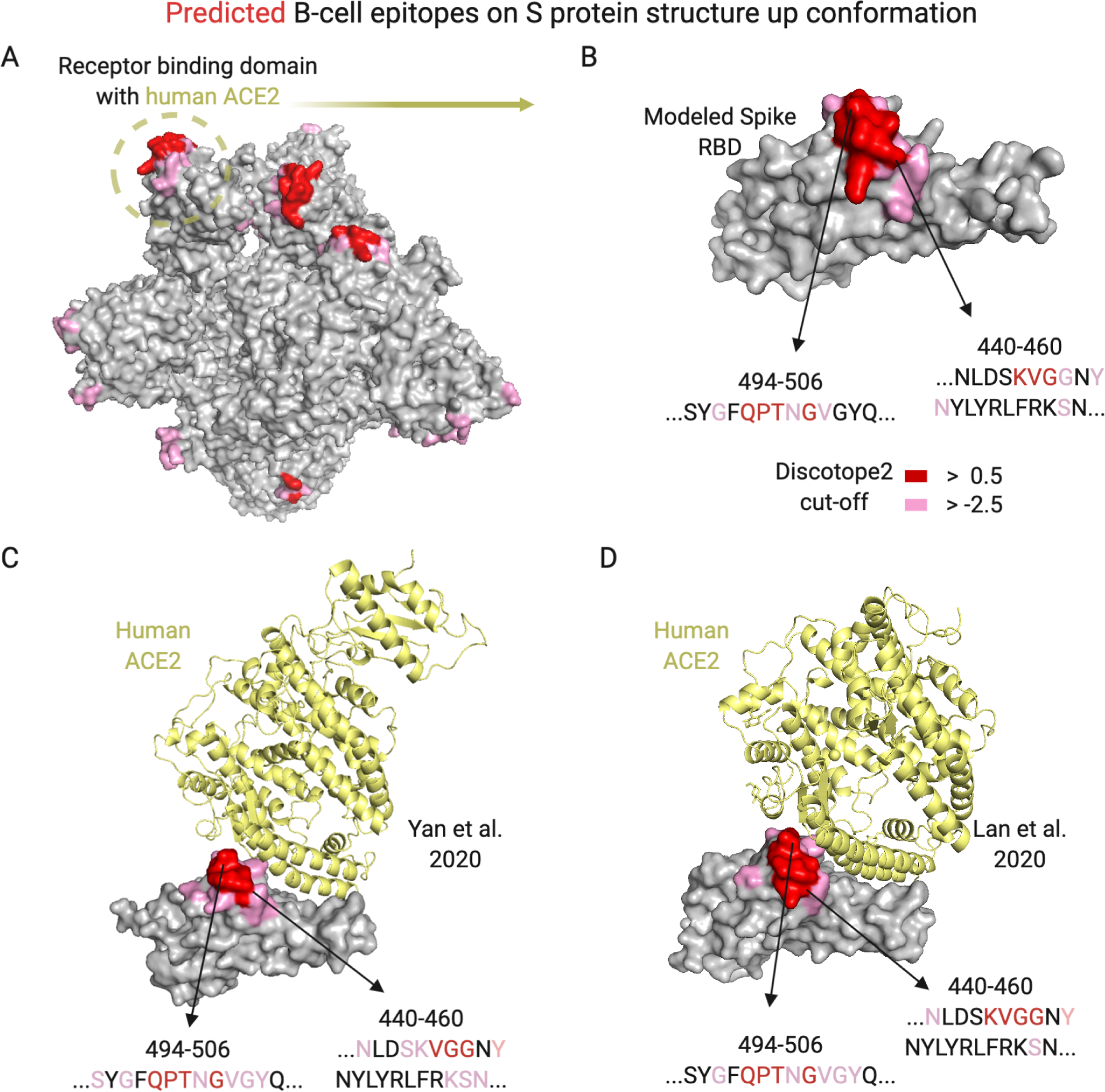
Predicted B-cell epitopes on 2019-nCoV spike (S) protein in up conformation. We scanned 3D structures of modeled (A) and experimentally solved (B) 2019-nCoV S protein in the up position with Discotope2 to assess potential antibody binding sites (B-cell epitopes). Red indicates residues with a score greater than 0.5 (with a cut-off for specificity of 90%), and pink indicates residues with score > −2.5 (with a cut-off for specificity of 80%). (**A**) Discotope2 identified multiple B-cell epitopes on the modeled 2019-nCov S protein. The top candidates cluster around the receptor binding domain (RBD). (**B**) The RBD B-cell epitotope cluster is comprised of two linear epitopes (440-460, 494-506). (**C**,**D**) Discotope2 identified highly similar B-cell epitopes with solved 2019-nCoV co-crystal structures from two independent research groups (19, 20). Predicted antibody binding sites overlap with the interacting surface for the known human entry receptor ACE2 (yellow).

For the 2019-nCoV S protein down conformation, Discotope2 identified a similar antibody binding site on the S protein potential RBD, but with fewer residues (17 residues, Fig. 5B). Residues 440-460 and 494-506 located at the top of the RBD show strong antibody binding properties (>0.5 raw scores). Discotope2 also identified another antibody binding site (residue 246-257) different from SARS-CoV but with lower scores (>-2.5 raw scores).

For the 2019-nCoV S protein up conformation, Discotope2 identified similar antibody binding sites compared to down conformation (440-460 and 494-506, Fig. 6). We wanted to explore whether these two linear epitopes are critical for viral interaction with the human entry protein ACE2. After our original analysis, two solved co-crystal structures of 2019-nCoV S protein RBD and human ACE2 become available (19, 20). We ran Discotope2 and obtained strikingly similar results (Fig. 6C and D). The main antibody binding site substantially overlaps with the interacting surface where ACE2 (yellow) binds to S protein, so an antibody binding to this surface is likely to block viral entry into cells.

Mutations in key antibody binding sites can help viruses evade the human immune system. We identified 8 point mutation sites on S protein from a cohort of 68 2019-nCoV samples (Fig. 5D). All mutations are distant from the S protein RBD, and one mutation (S247R) occurs near one of the predicted minor antibody binding site (residue 246-257).

In summary, we observed major structural and B-cell epitope similarity between SARS-CoV and 2019-nCoV spike proteins. RBDs in both proteins seem to be important B-cell epitopes for neutralizing antibodies, however, SARS-CoV appears to have larger attack surface than 2019-nCoV (Fig. 5A, 5B). Fortunately, we have not observed any mutations altering binding sites for this major B-cell epitope in 2019-nCoV.

### Promising candidates for 2019-nCoV vaccines

After understanding the landscape of T-cell and B-cell epitopes of 2019-nCoV, we summarize our results in Table 2 for promising candidates for 2019-nCoV vaccines. Ideal antigens should be presented by multiple MHC-I and MHC-II alleles in a general population and contain linear B-cell epitopes associated with neutralizing antibodies. Specifically, we first identified regions with high coverage of MHC-II, then ranked them based on their MHC-I coverage. We further search these top candidates with 90% sequence similarity on IEDB (33) to assess whether any candidate has been previously reported for being a B-cell epitope or T-cell epitope.

**Table 2.**
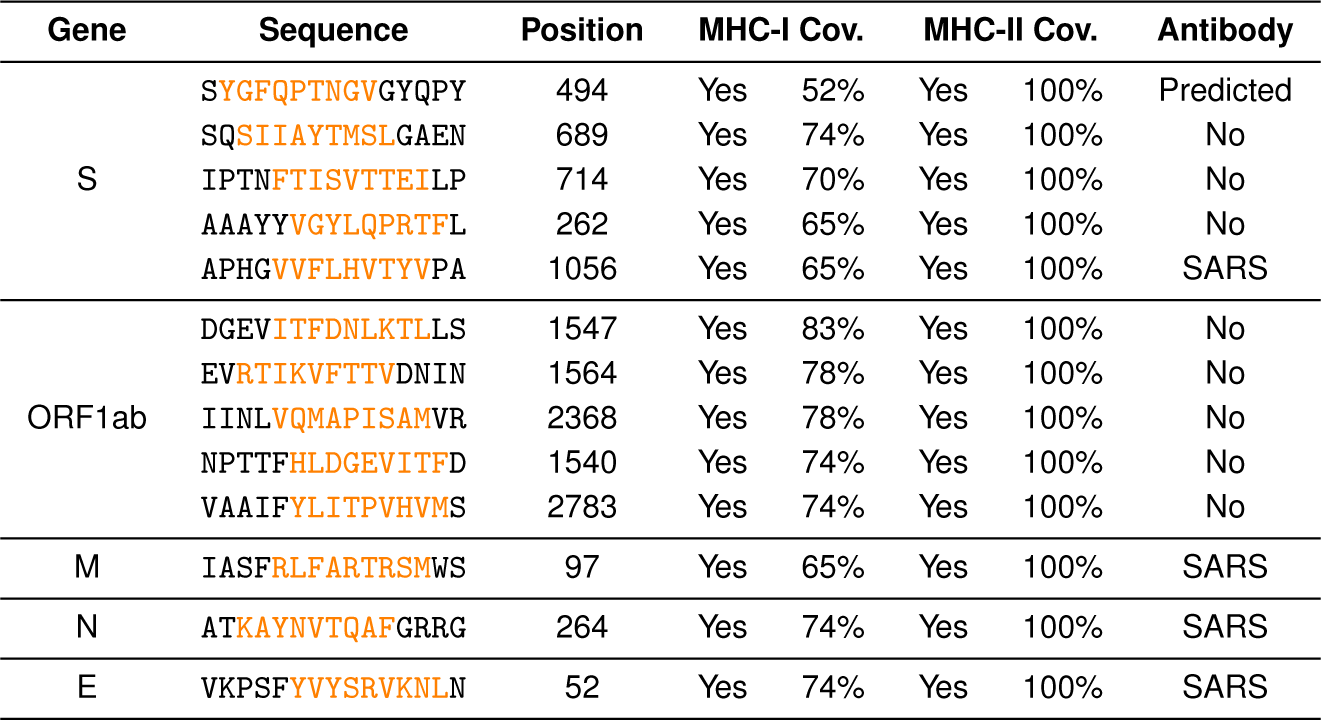
Top potential epitopes for key 2019-nCoV proteins. We ranked epitopes based on their likely coverage of presentation by MHC-I and MHC-II alleles. S protein 494-508 is highly ranked based on MHC presentation and is also one of the predicted top B-cell epitopes, localized near the S protein receptor binding domain (Fig. 5). MHC-I coverage is calculated by the 9mer with the highest MHC-I coverage for each epitope (highlighted in orange). All candidates are likely to be presented by both MHC-I and MHC-II. “SARS” under the antibody column indicates that one or more SARS homolog of this peptide is a known B-cell epitope.

Around 1000 peptide 15mers can be presented by a wide range of MHC-II (>33%), which is consistent with our knowledge about promiscuity of MHC-II presentation (15, 34). When further requiring a peptide to be presented by more than 33% or 66% of common MHC-I alleles, we narrowed the results to 405 or 44 promising T-cell epitopes, respectively (SI Appendix, Supplementary Tables 5 and 6). The top 13 candidates in key viral proteins can be presented by 52-83% of MHC-I alleles (Table 2). Most notably, S protein 494-508 is not only a good MHC presenter but also a predicted B-cell epitope near the S protein receptor binding domain (Fig. 5B), which indicates a good vaccine candidate.

When searching these 13 candidates on IEDB, we observed several of their homologous peptides in SARS are T-cell and B-cell epitopes. SARS peptide SIVAYTMSL is similar to 2019-nCoV peptide SQSIIAYMSLGAEN and elicits T-cell responses in chromium cytotoxicity assays (35). Previously discovered SARS B-cell epitopes LMSFPQAAPHGVVFLHV (36), ASFRLFARTRSMWSF(37), KRTATKQYNVTQAF-GRR (38), and SRVKNLNSSEGVPDLLV (39) share high homology with 2019-nCoV S protein 1056-1060, M 97-111, N 264-278, and E 52-66. These search results support validity of our computational pipeline.

## Discussion

In summary, we analyzed the 2019-nCoV viral genome for epitope candidates and found 405 likely T-Cell epitopes, with strong MHC-I and MHC-II presentation scores, as well as 2 potential neutralizing B-Cell epitopes on the S protein. We validated our methodology by running a similar analysis on SARS-CoV against known SARS T-cell and B-cell epitopes. Analyzing viral mutations across 68 patient samples we found that they have a tendency to occur in regions with strong MHC-I presentation, a possible form of immune evasion. However, no mutations are present near the spike protein receptor binding domain, which is critical for viral entrance to cells and neutralizing antibody binding.

We observed that significant portions of 2019-nCoV viral protein can be presented by multiple HLA-C alleles. This might be due to relatively conserved HLA-C binding motif on the 9th position (the main anchor residue) where Leucine, Methionine, Phenylalanine, Tyrosine are commonly favored across alleles (40). HLA-C*07:02, HLA-C*01:02, and HLA-C*06:02 all have >10% allele frequency in the Chinese population (41). This suggests the future epitope-based vaccines should include HLA-C related epitopes beyond the common HLA-A*02:01 and HLA-B*40:01 approach.

Similar to SARS-CoV (1, 8), the 2019-nCoV spike protein is likely to be immunogenic as it carries a number of both T-cell and B-cell epitopes. Recombinant surface protein vaccines have demonstrated strong clinical utilities in hepatitis B and human papillomavirus (HPV) vaccines (11, 42). A recombinant 2019-nCoV spike protein vaccine combined with other top epitopes and an appropriate adjuvant could be a reasonable next step for 2019-nCoV vaccine development. Recently, two biotech companies Moderna and Inovio have announced early clinical trials for spike protein based vaccines.

From an economic perspective, the development of coronavirus vaccines faces significant financial hurdles as most such viruses do not cause endemic infections. After an epidemic episode (e.g. SARS) there is little financial incentive for private companies to develop vaccines against a specific strain of virus (2, 8, 10). To address these incentive issues, governments might provide greater funding support to create vaccines against past and future outbreaks.

Our pipeline provides a framework to identify strong epitope-based vaccine candidates and might be applied against any unknown pathogens. We were able to model structures of 2019-nCoV spike protein as soon as the DNA sequence became available, and the modeled structure has high similarity with later solved crystal structures (SI Appendix, Fig. S1). This suggests the researchers can rely on homology remodeling when emerging pathogens first appear if the new pathogen shares high sequence similarity with an existing pathogen.

Previous animal studies show antigens with high MHC presentation scores are more likely to elicit strong T-cell responses, but the correlation between these responses and vaccine efficacy is relatively weak (7, 14, 43). When combined with future clinical data, our work can help the field untangle the relationship between antigen presentation and vaccine efficacy.

## Materials and Methods

We obtained 2019-nCoV (SARS-CoV-2019) and SARS-CoV reference sequence data from NCBI GeneBank (NC_045512 and NC_004718) (4, 16). We obtained viral sequences associated with 68 patients from GISAID on Feb 1st 2020.

We broke each gene sequence in 2019-nCoV into sliding windows and used NetMHCpan4 (17) and MARIA (15) to predict MHC-I and MHC-II presentation scores across 32 MHC alleles common in the Chinese population (41). We marked a peptide sequence as covered by MHC-I and MHC-II if it is presented by more then 33% of constituent alleles. For validation, we applied our methodology to known SARS T-cell epitopes and non-epitopes.

We predicted likely human antibody binding sites (B-cell epitopes) on SARS and 2019-nCoV S protein with Disctope2 (21) after first obtaining an approximate 3D structure of 2019-nCoV S protein by homology modeling SARS S protein (PDB: 6ACC) with SWISS-MODEL (31, 44). To validate our B-cell epitope predictions, we compared our top 3 B-cell epitopes for SARS S protein with previously experimentally identified epitopes (28–30).

For mutation identification, we translated viral genome sequences from patients into protein sequences and compared them with the reference sequence to identify mutations with edit distance analysis. We compared positions of point mutations with MHC-I or MHC-II presentable regions in the protein with Fisher’s exact test.

## ACKNOWLEDGMENTS

We thank Stanford Schor and Allen Lin for the discussion about coronaviruses. We thank all labs contributing to the global efforts of sequencing 2019-nCoV samples and we attached the full acknowledgment list in SI Appendix, Supplementary Table 1.

