## Supplementary Information/Appendix for "Potential T-cell and B-cell Epitopes of 2019-nCoV"

### Supporting Information Text

### Materials

**Viral sequences.** We obtained 2019-nCoV (SARS-CoV-2019) and SARS-CoV reference sequence data from NCBI GeneBank (NC_045512 and NC_004718) (1, 2). We then extracted the 2019-nCoV protein sequences of ORF1AB, S, ORF3A, E, M, ORF6, ORF7A, ORF7B, ORF8, N, and ORF10 based on the reference genome. We obtained viral sequences associated with 68 patients from GISAID on Feb 1st 2020 (Supplementary Table 1) (3). Submissions with a single viral gene are not included in the analysis. Full DNA and protein sequences of these 68 samples are available in Dataset S2 and S3.

**SARS epitopes.** We obtained known SARS T-cell and B-cell epitopes from the IEDB website (4).

### Methods

**MHC antigen presentation prediction.** We broke each gene sequence in 2019-nCoV into sliding windows of length 9, the median length of MHC-I ligands, and 15, the median length of MHC-II ligands. We used netMHCpan4 (5) and MARIA (6) to predict MHC-I and MHC-II presentation scores, respectively. We used 32 MHC alleles common in the Chinese population (>4% allele frequency, 7 HLA-A, 8 HLA-B, 9 HLA-C, 8 HLA-DRB1), as determined by an analysis of human populations (7). The complete list of common MHC alleles is included in Supplementary Table 2.

Both MARIA and netMHCpan4 return percentiles that characterize a peptide's likelihood of presentation relative to a preset distribution of random human peptide scores. Prior work recommends thresholds of 98% for NetMHCpan4 and 95% for MARIA to determine reasonable presenters. We also applied a more stringent 99.5% threshold for both NetMHCpan4 and MARIA. We used gene expression values of 50 TPM when running MARIA to reflect the high expression values of viral genes in human cells.

When aggregating alleles across MHC-I and MHC-II to report overall coverage, we marked a peptide sequence as covered if it is presented by more then 33% of common alleles. We chose 33% as a cut-off because it suggests a high (>90%) probability that at least one allele can present this peptide assuming the patient carries six MHC alleles (e.g. 2 As, 2 Bs, and 2 Cs) and the distribution of common MHC alleles in the population is random. We ranked potential T-cell epitopes based on their MHC-I and MHC-II presentation coverage across alleles.

**T-cell epitope validation.** We applied our methodology to known SARS T-cell epitopes and non-epitope SARS peptides to estimate our ability to predict 2019-nCoV epitopes. For MHC-I, we curated 17 experimentally determined HLA-A*02:01 associated CD8 T-cell epitopes and 1236 non-epitope 9mer sliding windows on SARS S protein (8–10). For MHC-II, we curated 3 experimentally determined CD4 T-cell epitopes and 246 non-epitopes on SARS S protein (11). No specific HLA-DR alleles were reported in the original study, so we used HLA-DRB1*09:01 and 15:01 (common alleles) to run MARIA. To calculate sensitivity and specificity for this validation set, we labeled any peptide sequence above the 98th (MHC-I) or 95th (MHC-II) percentile as a positive epitope prediction. Any sequence below that threshold we labeled as a negative prediction. We calculated AUC scores (AUROC) to estimate the overall performance of our methodology.

**Homology modeling of 2019-nCoV S protein.** We estimated the 3D structure of 2019-nCoV S protein by homology modeling SARS S protein with SWISS-MODEL (12). We modeled up and down conformations separately with two different SARS spike structures as templates (PDB: 6ACC for down and 6ACD for up) (13). S proteins from 2019-nCoV and SARS share 93% similarity. The modeled structures of 2019-nCoV S protein show high similarity with later Cryo-EM solved structures (Fig. S1, Datasets S4 and S5). Graphic rendering and analysis was performed with PyMOL (14).

**B-cell epitope prediction and validation.** We predicted likely human antibody binding sites (B-cell epitopes) on SARS and 2019-nCoV S protein with Disctope2 (15). Our analysis focused on neutralizing binding sites by only examining residues 1-600. The full prediction results can be found in Supplementary Tables 3 and 4. To validate our B-cell epitope predictions, we compared our top 3 B-cell epitopes for SARS S protein with previously experimentally identified epitopes from three independent studies (16–18).

**Mutation identification.** We aimed to determine whether there was any statistical relationship between regions of mutation and presentation. We translated viral genome sequences into protein sequences using BioPython (19) and indexes from the NCBI reference genome. 2019-nCoV nucleotide position 13468 contains a -1 frame shift signal and we adjusted accordingly to generate ORF1ab. We compared protein sequences from 68 patients to the reference sequence to identify mutations with edit distance analysis. Positions with poor quality reads (e.g. W or Y) were excluded from the analysis. The full sequence and mutation profiles can be found in Datasets S2 and S3. We compared positions of point mutations with MHC-I or MHC-II presentable regions in the protein with Fisher's exact test. Specifically, we use Fisher's exact test to compare the proportion of 9mers covered by >33% HLA-C alleles to the proportion covered by >33% of HLA-A and HLA-B alleles.

**Statistical analysis.** We computed Fisher's exact test with scipy (20) and AUROC with scikit-learn (21).

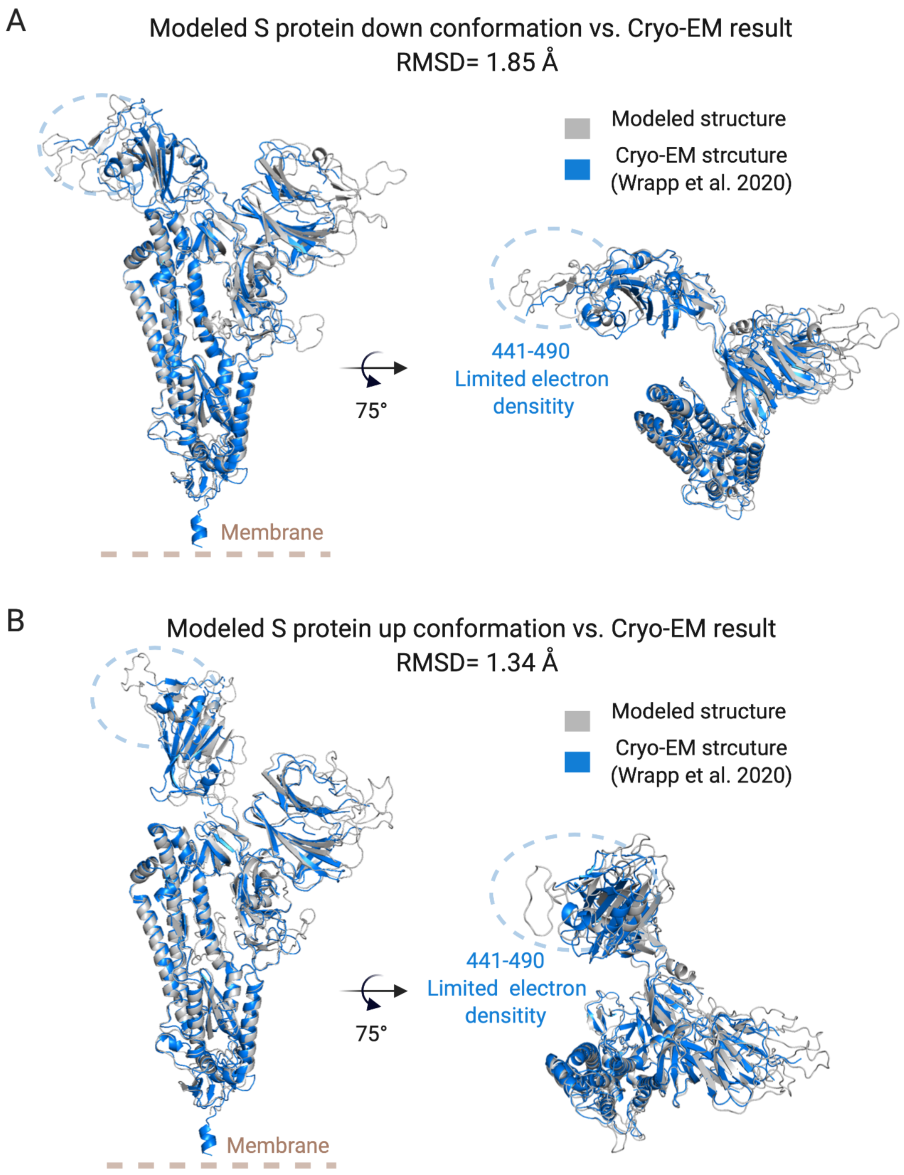


**Fig. S1. Comparison of modeled 2019-nCoV spike (S) protein structure and Cryo-EM solved structure**. 2019-nCoV S protein in both down (A) or up (B) conformations were homology modeled based on SARS-CoV S protein structures (PDB: 6ACC and 6ACD). Here we compare these modeled 3D structures to the Cryo-EM solved structure (PDB: 6VSD). Homology modeled structures achieved high similarity in both down and up conformations (RMSD = 1.85Å and 1.34Å). Notably, the flexible part (441-490) of the Cyro-EM structure has a high number of missing residues due to limited electron density, which limits our ability to perform B-cell epitope predictions.

### 1. Supplementary Tables and Files

**SI Dataset S1 (S1-supp-tables1-8.xlsx)**
An excel file containing several datasets and supplementary tables:

- **Supplementary Table 1**: Full details and acknowledgements for the viral genome data used
- **Supplementary Table 2**: Common HLA alleles among the Chinese population
- **Supplementary Table 3**: Discotope2 prediction for SARS-nCoV spike protein
- **Supplementary Table 4**: Discotope2 prediction for 2019-nCoV spike protein (modeled)
- **Supplementary Table 5**: NetMHCpan4 percentiles for all possible 9mer peptide sequences for 2019-nCoV protein
- **Supplementary Table 6**: MARIA percentiles for all possible 15mer peptide sequences for 2019-nCoV protein and peptides with high coverage for both MHC-I and MHC-II alleles
- **Supplementary Table 7**: Validation CD8 T-cell epitopes for SARS spike protein
- **Supplementary Table 8**: Validation CD4 T-cell epitopes for SARS spike protein

**SI Dataset S2 (S2-viral-dna-sequences.tsv)**
DNA sequences of 68 2019-nCoV samples.

**SI Dataset S3 (S3-viral-sequences-mutations.tsv)**
Protein sequence and mutation profiles of 68 2019-nCoV samples.

**SI Dataset S4 (S4-spike-model-down.pdb)**
PDB file for 2019-nCoV spike protein in down conformation via homology modeling.

**SI Dataset S5 (S5-spike-model-up.pdb)**
PDB file for 2019-nCoV spike protein in up conformation via homology modeling.
